# Genomic Evidence for Phototrophic Oxidation of Small Alkanes in a Member of the Chloroflexi Phylum

**DOI:** 10.1101/531582

**Authors:** Lewis M. Ward, Patrick M. Shih, James Hemp, Takeshi Kakegawa, Woodward W. Fischer, Shawn E. McGlynn

## Abstract

Recent genomic and microcosm based studies revealed a wide diversity of previously unknown microbial processes involved in alkane and methane metabolism. Here we described a new bacterial genome from a member of the Chloroflexi phylum—termed here *Candidatus* Chlorolinea photoalkanotrophicum—with cooccurring pathways for phototrophy and the oxidation of methane and/or other small alkanes. Recovered as a metagenome-assembled genome from microbial mats in an iron-rich hot spring in Japan, *Ca.* ‘C. photoalkanotrophicum’ forms a new lineage within the Chloroflexi phylum and expands the known metabolic diversity of this already diverse clade. *Ca.* ‘C. photoalkanotrophicum’ appears to be metabolically versatile, capable of phototrophy (via a Type 2 reaction center), aerobic respiration, nitrite reduction, oxidation of carbon monoxide, oxidation and incorporation of carbon from methane and/or other short-chain alkanes such as propane, and potentially carbon fixation via a novel pathway composed of hybridized components of the serine cycle and the 3-hydroxypropionate bi-cycle. The biochemical network of this organism is constructed from components from multiple organisms and pathways, further demonstrating the modular nature of metabolic machinery and the ecological and evolutionary importance of horizontal gene transfer in the establishment of novel pathways.

## Introduction

Microbial oxidation of alkanes facilitates the breakdown of environmental hydrocarbons including methane from geological, biological, and anthropogenic sources, and is therefore an important component of the global carbon cycle. Microbial fluxes of methane production and oxidation alone are on the order of one billion tons of methane per year (Reeburgh 2007). Understanding the diversity and activity of microbes that are involved in cycling of methane and other alkanes thus represents an important challenge to developing accurate climate models that are applicable both today for understanding global warming (e.g. Thauer 2010), and in the past, when methane may have played an important role in maintaining habitable conditions under a fainter sun deep in Earth history (e.g. Pavlov et al. 2000). Recently, awareness of archaeal metabolism of methane and other small alkanes has expanded dramatically, particularly through genome-centric metagenomics as well as enrichment-based approaches (e.g. Evans et al. 2015, Laso-Pérez et al. 2016, Vanwonterghem et al. 2016 Borrel et al. 2019, Chen et al. 2019); this spate of discoveries strongly suggests an incomplete understanding of the microbial cycling of small hydrocarbons that may be improved with further sampling of novel microbial diversity.

One hypothetical microbial metabolism is that of methane oxidation coupled to photosynthesis. This metabolism has previously been proposed (e.g. Vishniac 1960), and indirect evidence exists that is consistent with this metabolism playing a role in natural environments (e.g. environmental measurements of light-dependent methane oxidation, e.g. Oswald et al. 2015). A report exists of methane utilization by a phototrophic strain tentatively classified as *Rhodopseudomonas gelatinosa* (Wertlieb and Vishniac 1967), but to our knowledge was never confirmed; the strain in question is no longer available, and subsequent attempts to culture photomethanotrophic organisms have failed (e.g. Ratering 1996, Frantz 2009). The capacity for photomethanotrophy has therefore remained elusive. Here we describe a microbe with cooccurring pathways for phototrophy and the consumption of methane and/or other small alkanes such as propane, representing the first genomic evidence for a single organism with the potential for acquiring electrons and carbon from methane into a photosynthetic electron transport chain and biomass, respectively.

## Results

OHK40 was recovered as a metagenome-assembled genome (MAG) from shotgun metagenome sequencing at Okuoku-hachikurou Onsen (OHK) in Akita, Prefecture, Japan. This MAG consists of 301 contigs totaling 6.93 Mb and 5980 coding sequences. GC content is 67.8%, N50 is 36593. 47 RNAs were recovered. The genome was estimated by CheckM to be 98% complete, with 2.8% contamination based on presence/absence and redundancy of single-copy marker genes.

Multiple phylogenetic analyses each independently place OHK40 within the *Chloroflexaceae* clade of phototrophs in the Chloroflexi phylum (Figure 1). These analyses illustrate that OHK40 is most closely related to *Ca.* Chloroploca asiatica and *Ca.* Viridilinea mediisalina (Figure 1). Comparison of OHK40 to *Ca.* Chloroploca asiatica and *Ca*. Viridilinea mediisalina via OrthoANI identity (Yoon et al. 2017) was calculated to be ~71%, while the average amino acid identity (AAI) (Rodriguez and Konstantinidis 2014) was ~66%. A partial 16S sequence from OHK40 (348 nucleotides long) was determined to be 93% similarity to *Ca.* Viridilinea mediisalina. These pairwise differences suggest divergence of OHK from these other taxa to at least the genus level; classification with the Genome Taxonomy Database supports assignment of OHK40 to a novel genus within the Chloroflexaceae (Parks et al. 2018). The OHK40 genome is somewhat larger than that of its close relatives (~5.7 Mb for *Ca.* Chloroploca asiatica and *Ca.* Viridilinea mediisalina). However, the genome of OHK40 is still significantly smaller than the largest Chloroflexi genome known to date (8.7 Mb for *Kouleothrix aurantiaca*, 26). The larger genome size of OHK40 is associated with 160 annotated coding sequences that were not recovered in the draft genomes of either of the closely related species *Ca.* Chloroploca asiatica or *Ca*. Viridilinea mediisalina (Supplemental Table 1-3)—these represent candidates for recent HGT into OHK40. Protein phylogenies were used to verify HGT into OHK40 of genes coding for metabolically relevant proteins as discussed below (Figure 2, Supplemental Figures 1-6).

**Figure 1:**
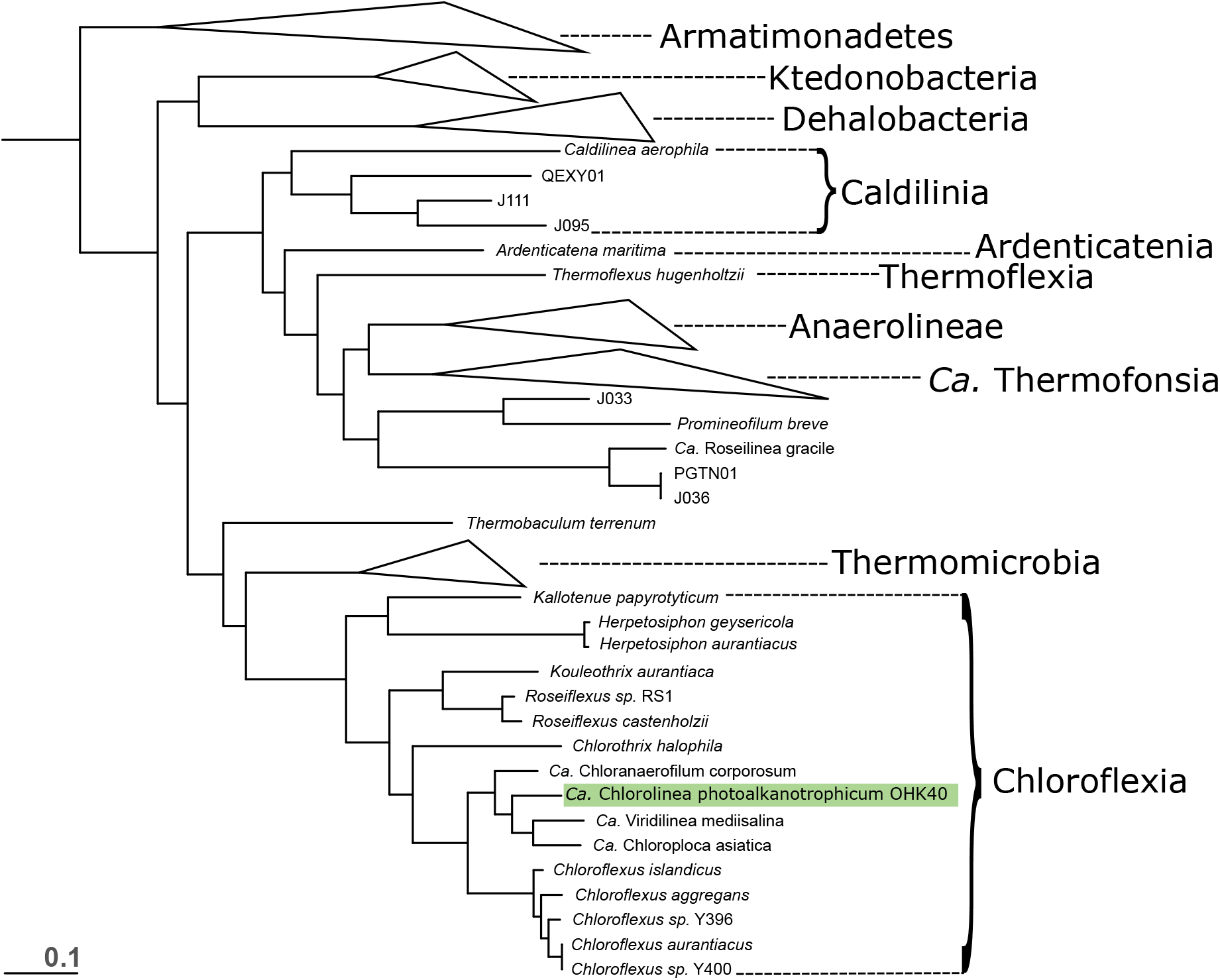
Phylogeny of the Chloroflexi phylum, built with concatenated ribosomal protein sequences following methods from Hug et al. 2016. The analysis contains members of the Chloroflexi phylum previously described and members of the closely related phylum Armatimonadetes as an outgroup. All nodes recovered TBE support values greater than 0.7. In cases where reference genomes have a unique strain name or identifier, this was included; otherwise Genbank WGS genome prefixes were used.

**Figure 2:**
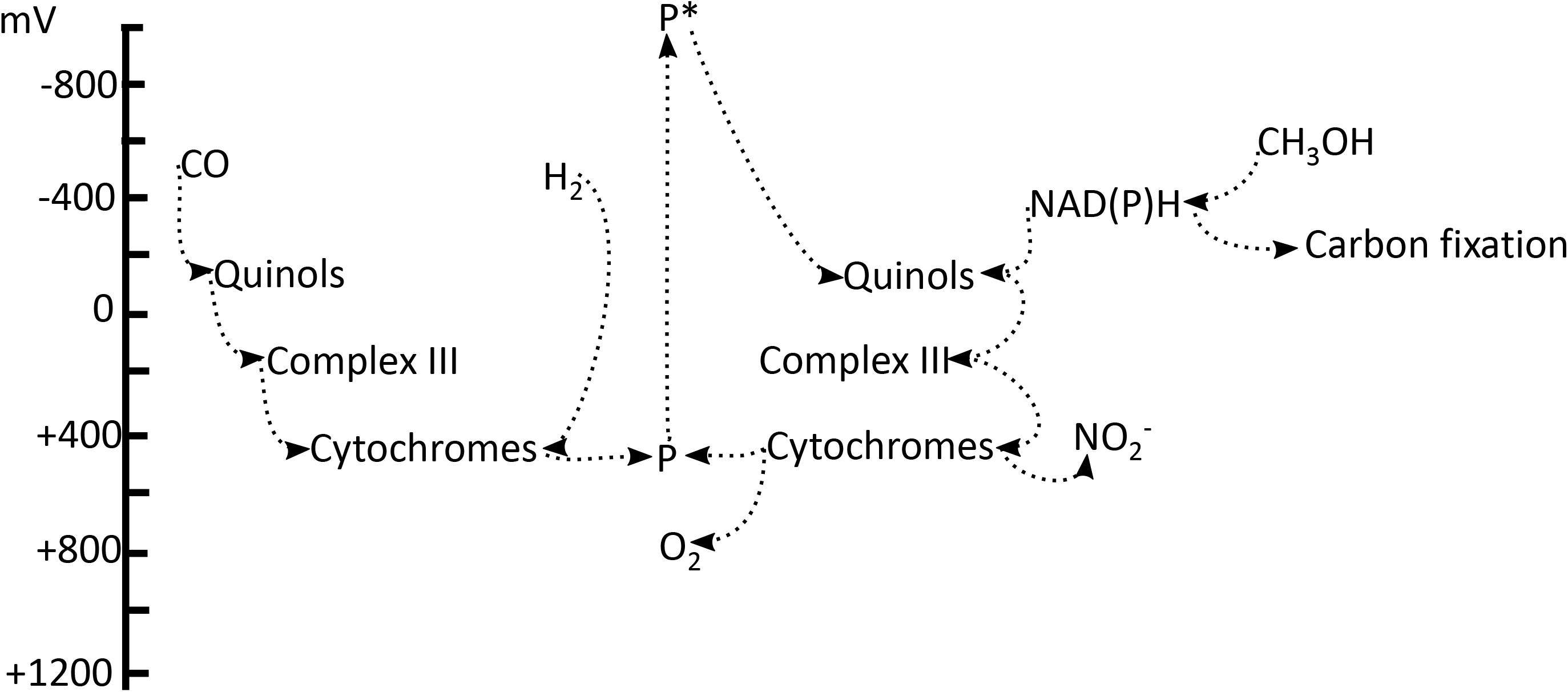
Phylogeny and structure of the methane monooxygenase hydroxylase (MMOH) from the Ca. C photoalkanotrophicum genome. A) Phylogeny of the protein sequence of the alpha chain of the A subunit of soluble methane monooxygenase (SmmoA), showing position of OHK40 relative to other sequences available from NCBI Genbank and WGS databases, on a branch near members of the genus *Sulfobacillus* and the uncultured gammaproteobacterial lineage UBA981, suggesting that OHK40 acquired this enzyme via horizontal gene transfer from a donor outside the Chloroflexi phylum. B) Structural overlay of the modeled structure (yellow) with pdb entry 1MTY (MMOH) chain D (MmoX) from *Methylococcus capsulatus* (Bath) (blue). C) Expanded view of the di-iron active site of the pdb derived structure 1MTY with the model (yellow). Nitrogen appears as a blue ball, oxygen in red, iron in rust.

Like closely related members of the Chloroflexaceae, the OHK40 genome encodes pathways for aerobic respiration (via an A-family and a B-family heme copper oxidoreductase), phototrophy via a Type 2 (quinone-reducing) reaction center, an alternative complex III, and synthesis of bacteriochlorophylls *a* and *c* (e.g. BchH, BchD, BchI, BchM, AcsF, BchL, BchN, BchB, BchX, BchY, BchZ, BchF, BchG, BchQ, BchU, and BchK). Additionally, OHK40 encodes a soluble methane monooxygenase (sMMO) (including all subunits found in closely related sequences—the alpha and beta chains of the A subunit (hydroxylase), component C (reductase), and regulatory protein B—but lacks the gamma chain of the A subunit typical of sMMO complexes, Rosenzweig et al. 1997). These proteins were recovered on a fairly long (~68kb) contig together with many hypothetical and annotated metabolic genes (e.g. sulfate permease, mannonate dehydratase, the starvation sensing protein RspA, and multiple peptide ABC transporters) that were determined via similarity and phylogenetic analyses to be most closely related to those from members of the Chloroflexaceae and therefore to have been vertically inherited genes that belong in the OHK40 genome, strengthening interpretations that this contig belongs in the OHK40 genome and was not recovered due to contamination of the MAG.A cytochrome *c*_552_ may enable nitrite reduction to ammonium, and a Form I CO oxidoreductase suggests a capacity to oxidize CO. The genome encodes a partial 3-hydroxypropionate cycle that is potentially linked to components of the serine cycle, in which carboxylation is performed by the left branch of the 3-hydroxypropionate bi-cycle (3HP) while glyoxylate produced as a byproduct of the 3HP cycle and methane- or propane-derived carbon are incorporated via the serine cycle (Figure 3, and described in detail below).

**Figure 3:**
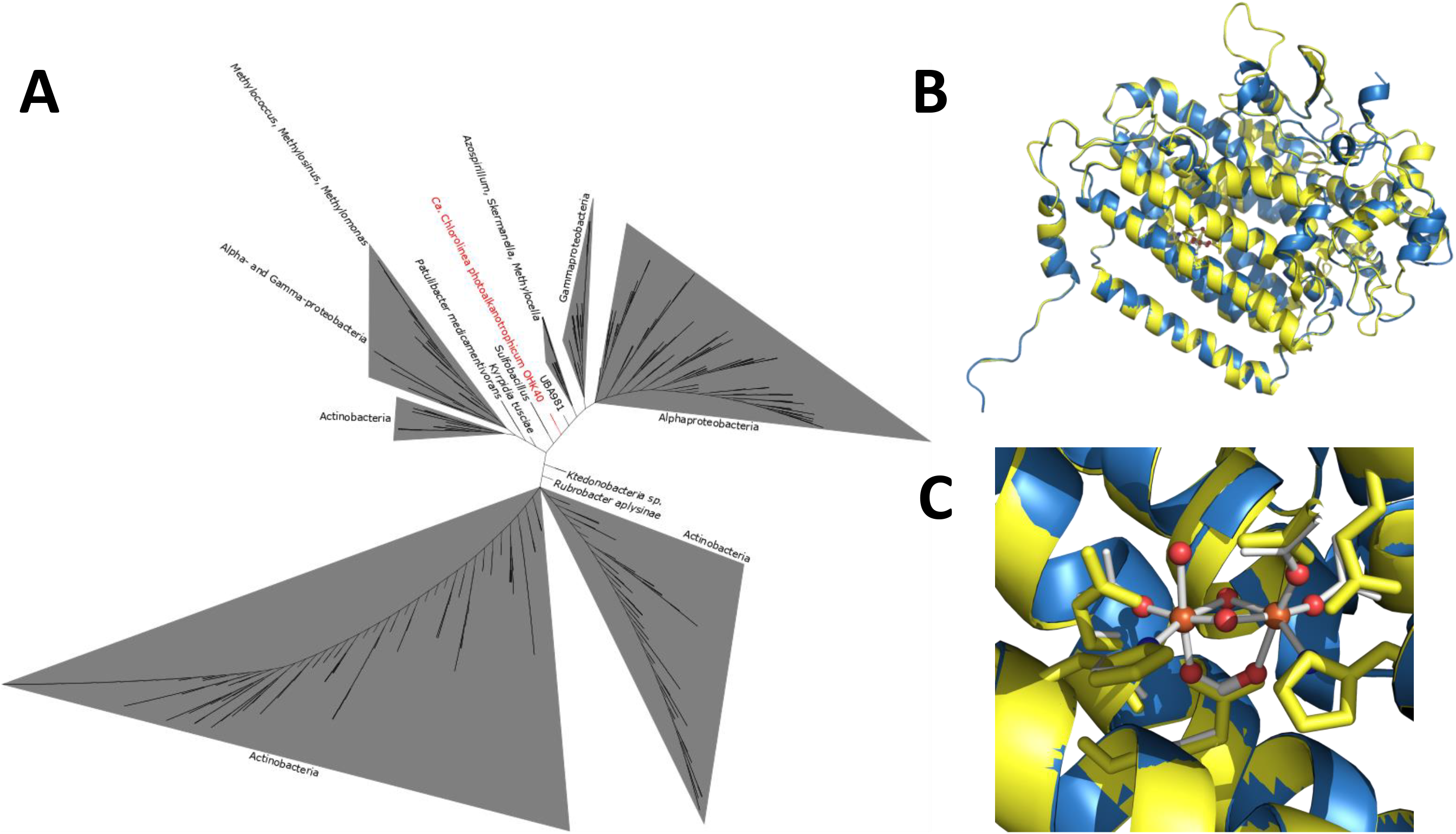
Diagram of putative carbon metabolism in OHK40, including incorporation of methane and bicarbonate into organic carbon via components of the serine and 3HP cycles, respectively. Dotted arrows indicate steps which were not recovered in the MAG, but which would enable a more complete hybridized pathway as discussed in the text. Red arrows indicate steps that appear to have been acquired in OHK via HGT since divergence from *Ca.* Chloroploca asiatica and *Ca*. Virdilinea mediisalina. A dotted blue arrows indicate a step which is not encoded by proteins annotated to perform this function, but for which close homologs that could potentially perform this step are encoded (e.g. acetolactate synthase for tartronate-semialdehyde synthase). Stoichiometry is 1:1 products to reactants for all steps, with the exception of step 15, which takes 2 glyoxylate as input and produces one tartronic semialdehyde and one CO_2_. 1) methane/propane monooxygenase; 2) alcohol dehydrogenase; 3) formaldehyde dehydrogenase; 4) formate dehydrogenase; 5) serine hydroxymethyltransferase; 6) serine deaminase; 7) serine glyoxylate aminotransferase; 8) hydroxypyruvate reductase; 9) glycerate kinase; 10) enolase; 11) phosphoenolpyruvate carboxylase; 12) malate dehydrogenase; 13) 2-hydroxy-3-oxopropionate reductase; 14) hydroxypyruvate isomerase; 15) tartronate-semialdehyde synthase; 16) malyl-CoA lyase; 17) acetyl-CoA carboxylase; 18) malonyl-CoA reductase; 19) propionyl-CoA synthase; 20) propionyl-CoA carboxylase; 21) methylmalonyl-CoA epimerase; 22) methylmalonyl-CoA mutase; 23) succinyl-CoA:(S)-malate-CoA transferase; 24) succinate dehydrogenase; 25) fumarate hydratase; 26) acetone monooxygenase; 27) methyl acetate hydrolase.

OHK40 does not encode genes for nitrogen fixation, canonical denitrification (i.e. the stepwise reduction of NO_3_^−^ to NO_2_^−^, NO, N_2_O, and finally N_2_,), the RuMP pathway for methane incorporation, or known genes for dissimilatory oxidation or reduction of sulfur- or iron-bearing compounds. The OHK40 genome does not encode the catalytic subunits of an uptake hydrogenase (HypA, HypC-F); however the genome does have assembly proteins for a NiFe uptake hydrogenase homologous to those from other phototrophic Chloroflexi (e.g. Klatt et al. 2013). These genes are at the end of a contig and the portion of the genome corresponding to that of *Chloroflexus aggregans* and *Roseiflexus castenholzii* that encodes catalytic subunit genes of the hydrogenase are missing in the OHK40 genome, suggesting that these genes may be encoded in the source genome but were not recovered in the MAG (despite the low MetaPOAP False Negative estimate ~0.02).

Protein phylogenies for multiple phototrophy, respiration, and carbon fixation genes are congruent with organismal phylogenies within the Chloroflexaceae (Supplemental Figures 1-3), consistent with vertical inheritance of these traits from the last common ancestor of the clade. In contrast, the putative sMMO and numerous other proteins (e.g. cytochrome *c*_552_ nitrite reductase, CO dehydrogenase, and methyl acetate hydroxylase) appear to have been acquired via HGT from more distantly related taxa (Figure 2, Supplemental Figures 5-10). The putative sMMO protein sequence in OHK40 is most closely related to sequences from uncultured organisms including the putatively methanotrophic gammaproteobacterium UB981 on a branch of the multisubunit monooxgenase tree between verified sMMO proteins from obligate methanotrophic Proteobacteria and a clade that includes the propane monooxygenase (PrMO) of *Methylocella sylvestris* (Crombie and Murrell 2014) (Figure 2). Moreover, this enzyme family typically oxidizes a broad range of substrates including methane, propane, and other small hydrocarbons *in vitro*, whether or not this allows growth on a range of substrates *in vivo* (Colby et al. 1977, Coleman et al. 2006, Hakemian and Rosenzweig 2007). Interpretations of the full range and preferred substrate(s) of the monooxygenase in OHK40 are therefore tentative pending experimental data following isolation or enrichment of this organism.

## Discussion and conclusions

The metabolic coupling of phototrophy and methanotrophy (photomethanotrophy) has been hypothesized to be viable for decades (e.g. Vishniac 1960), but neither this capacity nor the more general phototrophic oxidation of small alkanes (photoalkanotrophy) has never previously been confirmed to exist in a single organism. The genome described here, OHK40, provides the first genomic description of the potential for photomethanotrophy. Given the degree of genetic divergence of OHK40 and its apparent unique metabolic attributes, we propose a new genus and species designation within the Chloroflexaceae family of Chloroflexi, *Candidatus* Chlorolinea photoalkanotrophicum, pending isolation and further characterization. The designation of OHK40 as a new genus-level lineage is consistent with recent proposals for standardized genome sequence-based taxonomy (Parks et al. 2018).

The coupling of methanotrophy or propanotrophy to phototrophy would be enabled by the modular nature of high-potential electron transfer pathways (Ward et al. 2018a, Fischer et al. 2016, Shih et al. 2017). Electrons derived from oxidation of single-carbon compounds or short alkanes can be fed into the phototrophic reaction center to drive cyclic electron flow for energy conservation and subsequently used for carbon fixation (Figure 4). Carbon could also be directly incorporated into biomass from methane or propane via methanol and formaldehyde in order to supplement or replace the more energetically costly fixation of dissolved inorganic carbon (DIC, e.g. CO_2_ and HCO_3_^−^) (Figure 3) (see below).

**Figure 4:**
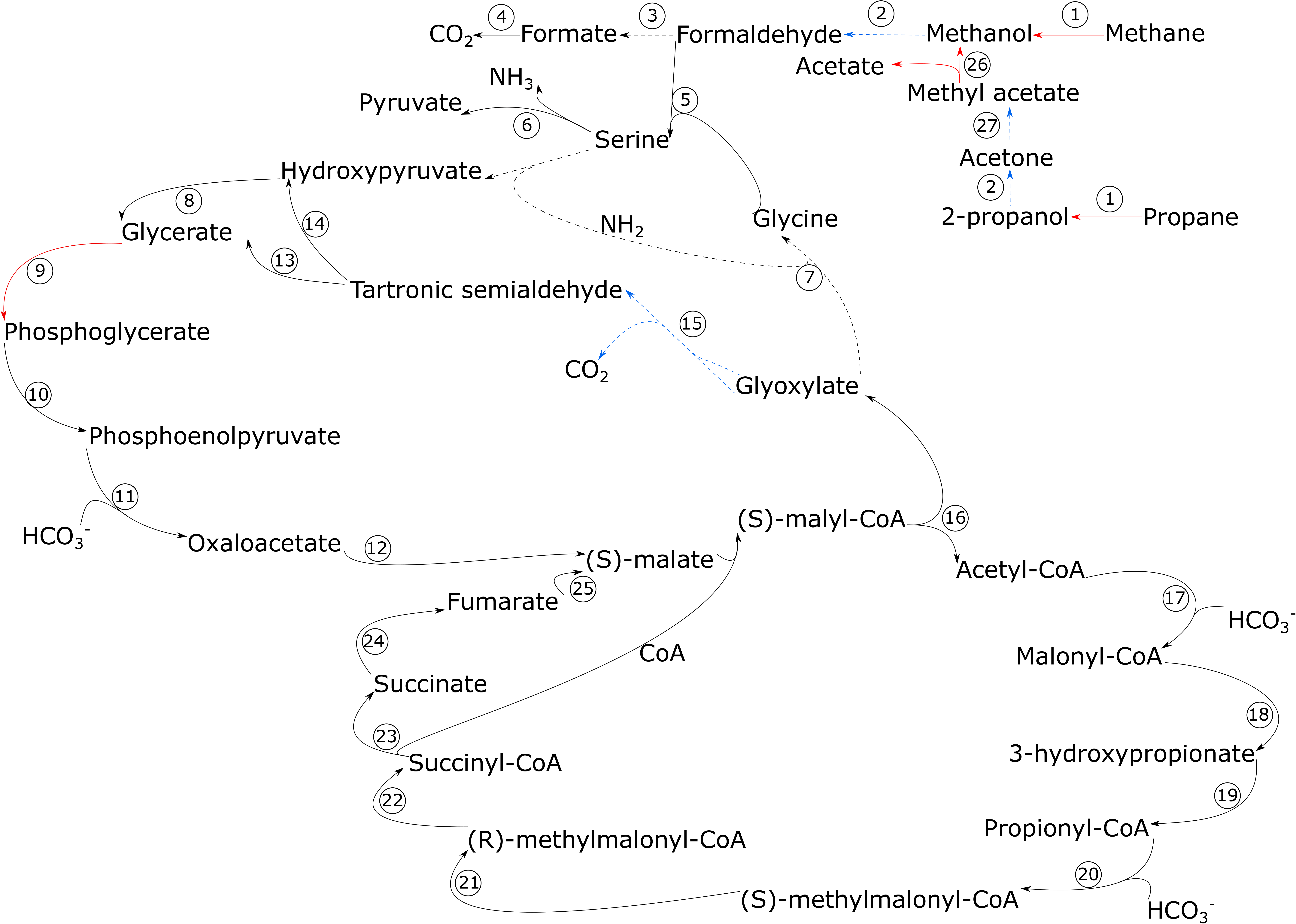
Diagram of putative electron transfer in OHK40 in redox potential space. Electrons sourced from methanol (derived from oxidation of methane or propane), carbon monoxide, or other donors are siphoned into the phototrophic electron transfer chain for conservation of energy (i.e. buildup of proton motive force) before being transferred uphill to reduced electron carriers such as NAD(P)H for carbon fixation. This is in contrast to more oxidized electron donors for photosynthesis such as H_2_O or NO_2_^−^ which must be fed directly into the reaction center (e.g. Fischer et al. 2016).

In nonphototrophic aerobic methanotrophs, electrons from methane are run through electron transport chains and ultimately donated to O_2_ (or, rarely, oxidized nitrogen species, e.g. Skennerton et al. 2015) at complex IV; this respiration of methane-derived electrons results in a small, finite number of protons pumped across the membrane per methane molecule oxidized, in contrast to phototrophs which can conserve energy from light via cyclic electron transfer without net consumption of electrons. As a result, the energetic yield per methane molecule of photomethanotrophy could be much higher than purely respiratory methanotrophy.

The 3-hydroxypropionate bi-cycle is the characteristic carbon fixation pathway found in most phototrophic members of the Chloroflexaceae (Ward et al. 2018a, Shih et al. 2017). *Ca.* C. photoalkanotrophicum encodes portions of the 3-hydroxypropionate bi-cycle, but is lacking some key proteins (Figure 3). The canonical 3HP pathways involves two cycles for carboxylation and the subsequent incorporation of glyoxylate. *Ca.* C. photoalkanotrophicum does not encode the second (right) cycle of 3HP which is responsible for conversion of glyoxylate into pyruvate. This suggests that *Ca.* C. photoalkanotrophicum may primarily function as a photoheterotroph. It is possible, however, that *Ca.* C. photoalkanotrophicum may encode an alternative mechanism of glyoxylate incorporation via conversion to glycerate by way of components of the serine cycle. This proposed carbon metabolism is expected to function similarly whether *Ca.* C. photoalkanotrophicum is consuming methane or propane, as these compounds share an uptake pathway via methanol as an early intermediate (Figure 3).

*Ca.* C. photoalkanotrophicum recovered nearly the complete set of genes involved in the proposed hybrid 3HP/serine cycle. The only step that was not recovered was serine/glyoxylate aminotransferase (step 7 in Figure 2). Although not recovered in the MAG, the presence in the genome of a gene encoding serine-glyoxylate aminotransferase would enable a complete hybridized 3HP/serine cycle for autotrophic carbon fixation through either or both bicarbonate and alkanes such as methane and propane. Its absence would result in a cycle which could incorporate methane or other short alkanes into some, but not all, biomolecules, with the remainder coming from exogenous organic carbon (mixotrophy) or a modified form of the 3HP cycle (autotrophy). In this case, methane and propane would only be directly incorporated into serine-derived biomass, including cysteine, glycine, threonine, and porphyrins such as bacteriochlorophylls (themselves derived primarily from glycine). The remainder of biomass could be produced through the 3HP variant present in *Ca.* C. photoalkanotrophicum, or could be derived from exogenous organic carbon leading to a mixotrophic lifestyle (which is common among phototrophic Chloroflexi, e.g. Klatt et al. 2013). MetaPOAP analyses show that the probability of failure to recover this one gene in the MAG is low (~0.02), but considered in the context of the entire 3HP/serine cycle, the probability of failing to recover at least one gene out of the ~20 involved in the total pathway is fairly high (~0.33, assuming failure to recover any one gene is ~0.02). It is therefore difficult to reject the hypothesis that *Ca.* C. photoalkanotrophicum may encode a complete 3HP/serine cycle. There is already precedence for the plasticity of Chloroflexi to transition between photoautotrophy and photoheterotrophy and to mix and match various metabolic pathways within central carbon metabolism, as various members have already been demonstrated to have lost enzymes involved in the 3HP pathway with some simultaneously acquiring other carbon fixation pathways (i.e., the Calvin cycle via RuBisCO and phosphoribulokinase) (Ward et al. 2018a, Shih et al. 2017).

Additionally, it appears that *Ca.* C. photoalkanotrophicum could be capable of harvesting electrons from carbon monoxide via a Form 1 CO dehydrogenase to feed into the phototrophy pathway. During carbon monoxide metabolism, CO is oxidized completely to CO_2_; CO-derived carbon cannot be directly incorporated into biomass, but electrons from CO could be used to fix DIC into biomass. To our knowledge, this is the first evidence for phototrophic CO metabolism in the Chloroflexi. Previous reports of CO metabolism in Chloroflexi have been restricted to nonphototrophic lineages (e.g. Islam et al. 2019). Phototrophic CO oxidation has previously been proposed for the anoxygenic phototrophic proteobacterium *Rhodopseudomonas palustris* (Larimer et al. 2004). A separate route for electron intake could occur by the activity of a [NiFe] hydrogenase, which could feed electrons to the phototrophic reaction center and ultimately to CO_2_ for carbon fixation or onto a respiratory electron acceptor (Figure 4).

The discovery of alkanotrophy generally and perhaps methanotrophy specifically in a member of the Chloroflexi phylum expands the known metabolic diversity of this phylum and further reinforces interpretations of the Chloroflexi as one of the most metabolically diverse bacterial phyla, following recent descriptions of Chloroflexi with diverse metabolic traits including iron reduction (Kawaichi et al. 2013), complete denitrification (Hemp et al. 2015c), sulfate reduction (Anantharaman et al. 2018), nitrite oxidation (Sorokin et al. 2012), and lithoautotrophic hydrogen oxidation (Ward et al. 2018f). The discovery of putative photomethanotrophy helps to fill a gap in thermodynamically favorable metabolisms hypothesized to exist long before their discovery in the environment, alongside metabolisms such as anammox, comammox, and photoferrotrophy (Kuenen 2008, Daims et al. 2015, van Kessel et al. 2015, Widdel et al. 1993, Vishniac 1960).

While *Ca.* C. photoalkanotrophicum is the first described genome of a putatively photomethanotrophic organism, this metabolism may be more broadly distributed. Though to our knowledge it has never been discussed in the literature, genes for methanotrophy and phototrophy also cooccur in several sequenced members of the Proteobacteria including *Methylocella silvestris* (NCBI-WP_012591068.1), *Methylocystis palsarum* (NCBI-WP_091684863.1), and *Methylocystis rosea* (NCBI-WP_018406831.1), though their capacity for photomethanotrophy has not yet been demonstrated. These organisms are not closely related to *Ca.* C. photoalkanotrophicum, and so photomethanotrophy may have evolved convergently in both the Chloroflexi and Proteobacteria phyla. If photomethanotrophy is more widespread, it may play a previously unrecognized role in modulating methane release in methane-rich photic environments such as wetlands, stratified lakes, and thawing permafrost. Photomethanotrophy may even have contributed to previous observations of light-dependent methane oxidation in diverse environments (e.g. Oswald et al. 2015). However, the derived phylogenetic placement of putative photomethanotrophs and the obligate O_2_ requirement of aerobic photomethanotrophy suggest that this metabolism did not play a role early in Earth history, before the rise of oxygen (supplemental information).

## Materials and methods

### Geological context and sample collection

The metagenome-assembled genome described here was derived from shotgun metagenomic sequencing of microbial communities of Okuoku-hachikurou Onsen (OHK) in Akita, Prefecture, Japan. OHK is an iron-carbonate hot spring (Takashima et al. 2011, Ward et al. 2017a). This spring is remarkable for its unique iron-oxidizing microbial community and the accumulation of iron-rich tufa (authigenic mineral cement) that has some textural features in common with sedimentary iron formations deposited during Precambrian time (Ward et al. 2017a). In brief, the geochemistry of OHK derives from source waters supersaturated in CO_2_, anoxic, pH 6.8, ~45 °C, and containing ~22 μM dissolved NH_3_/NH_4_^+^ and ~114 μM dissolved Fe^2+^ (Ward et al. 2017a).

Samples for shotgun metagenomic sequencing were collected in September 2016 from the “Shallow Source” and “Canal” sites described in (Ward et al. 2017a). Thin biofilms (<1 mm) were scraped from mineral precipitates using sterile forceps and spatulas (~0.25 cm^3^ of material). Cells were lysed and DNA preserved in the field using a Zymo Terralyzer BashingBead Matrix and Xpedition Lysis Buffer. Cells were disrupted immediately by attaching tubes to the blade of a cordless reciprocating saw, which was run for 60 s.

### Sequencing and analysis

Metagenomic sequencing and analysis, including genome binning, followed methods described previously (Ward et al. 2018a, Ward et al. 2018f) and described in the SI.

Presence of metabolic pathways of interest was predicted with MetaPOAP (Ward et al. 2018b) to check for False Positives (contamination) or False Negatives (genes present in source genome but not recovered in MAG). Phylogenetic trees were calculated using RAxML Stamatakis 2014) on the Cipres science gateway (Miller et al. 2010). Transfer bootstrap support values were calculated by BOOSTER (Lemoine et al. 2018), and trees were visualized with the Interactive Tree of Life viewer (Letunic and Bork 2016). Taxonomic assignment of the OHK40 genome was determined using markers including placement in reference phylogenies built using the RpoB protein (a single copy marker that is typically vertically inherited and more commonly recovered in MAGs than 16S rDNA, e.g. Ward et al. 2018a) and concatenated ribosomal protein sequences (following methods from Hug et al. 2016) and using a partial 16S rDNA sequence recovered in the genome. Taxonomic assignment was further confirmed with GTDB-Tk (Parks et al. 2018).

Protein structural modeling of the methane monooxygenase was done with the SWISS-MODEL workspace (Bienert et al. 2017). The predicted hydroxylase SMMO subunit was structurally aligned to the protein data base structure 1MTY within PyMOL.

## Acknowledgements

LMW acknowledges support from NASA NESSF (#NNX16AP39H), NSF (#OISE 1639454), NSF GROW (#DGE 1144469), the Earth-Life Science Institute Origins Network (EON), and the Agouron Institute. P.M.S. was supported by The Branco Weiss Fellowship - Society in Science from ETH Zurich. WWF acknowledges the generous support of the Caltech Center for Environment Microbe Interactions, NASA Exobiology (#NNX16AJ57G), and the Simons Foundation Collaboration on the Origins of Life (SCOL). SEM is supported by NSF Award 1724300, JSPS KAKENHI Grant Number 18H01325, and the Astrobiology Center Program of the National Institutes of Natural Sciences (grant no. AB311013).

## Supplemental Information

### Supplemental Methods

Upon return to the lab, microbial DNA was extracted and purified with a Zymo Soil/Fecal DNA extraction kit. Following extraction, DNA was quantified with a Qubit 3.0 fluorimeter (Life Technologies, Carlsbad, CA) according to manufacturer’s instructions. Purified DNA was submitted to SeqMatic LLC (Fremont, CA) for library preparation and 2×100 bp paired-end sequencing via Illumina HiSeq 4000 technology. The “Shallow Source” and “Canal” samples were prepared as separate libraries and multiplexed in a single sequencing lane with one sample from another project (Ward 2017). Raw sequence reads from the two samples were coassembled with MegaHit v. 1.02 (Li et l. 2016). Genome bins were constructed using MetaBAT (Kang et al. 2015), CONCOCT (Alneberg et al. 2013), and MaxBin (Wu et al. 2014) before being dereplicated and refined with DASTool (Sieber et al. 2018). Genome bins were assessed for completeness and contamination using CheckM (Parks et al. 2015) and contamination reduced with RefineM (Parks et al. 2017). The OHK40 genome was uploaded to RAST for preliminary annotation and characterization (Aziz et al. 2008). Sequences of ribosomal and metabolic proteins used in analyses (see below) were identified locally with the *tblastn* function of BLAST+ (Camacho et al. 2008), aligned with MUSCLE (68), and manually curated in Jalview (Waterhouse et al. 2009). Positive BLAST hits were considered to be full length (e.g. >90% the shortest reference sequence from an isolate genome) with *e*-values greater than 1e-20. Genes of interest were confirmed to be located on large, well-assembled (i.e. 10s of kb) contigs and screened against outlier (e.g. likely contaminant) contigs as determined by CheckM (Parks et al. 2015) and RefineM (Parks et al. 2017) using tetranucleotide, GC, and coding density content. To further confirm that genes of interest belong in the OHK40 genome and were not due to contamination, conserved marker genes (e.g. encoding ribosomal and central carbon metabolism proteins) were identified collocated on contigs with genes of interest and BLAST searches were performed against the Genbank database to confirm that these markers belong to a member of the Chloroflexaceae (i.e. belong to the OHK40 genome, and are not contamination).

### Supplemental Discussion

While soluble methane monooxygenase (sMMO) was classically considered to only occur along with particulate methane monooxygenase (pMMO) in aerobic methanotrophs (e.g. 10), more recent work has revealed that organisms encoding sMMO but not pMMO are not uncommon, especially in the case of facultative methanotrophs which may have acquired the capacity for methanotrophy via HGT (Semrau et al. 2011).

Among remaining undiscovered metabolisms, phototrophic ammonia oxidation (“photoammox”) is a viable metabolism that could be biochemically wired in a fashion similar to photomethanotrophy as described here: O_2_-enabled (via ammonia monooxygenase, AMO) anoxygenic phototrophic ammonia oxidation. AMO is homologous to particulate methane monooxygenase, and could work similarly to methane monooxygenase to activate ammonia using O_2_, producing hydroxylamine. Hydroxylamine oxidase would then produce nitrite and yield biologically useful electrons that could by fed into the phototrophic electron transport chain in a manner analogous to the photomethanotrophy pathway described here. No organism has yet been described which encodes both a phototrophic reaction center and genes for ammonia oxidation, though phototrophic nitrite oxidation by members of the Proteobacteria has recently been described (Hemp et al. 2016).

Methane oxidizing bacteria typically oxidize ammonia to nitrite due to the promiscuity of methane monooxygenase for ammonia and several other substrates including short hydrocarbons (Hanson and Hanson 1996), and so *Ca.* C. photoalkanotrophicum may be responsible for a minor, incidental amount of phototrophic ammonia cycling. The initial oxidation of ammonia by sMMO would produce hydroxylamine (e.g. Hanson and Hanson 1996). Hydroxylamine could rapidly react with dissolved Fe^2+^ found in the environment from which *Ca.* C. photoalkanotrophicum was recovered (Ward et al. 2017a), but this organism also encodes genes for cytochrome c552 nitrite reductase which is capable of reducing hydroxylamine to ammonia (Einsle et al. 1999); genes for this enzyme appear to have been acquired via HGT, and may be an adaptation to avoid hydroxylamine toxicity due to incidental ammonia oxidation—a strategy observed in some denitrifying methanotrophs (e.g. Skennerton et al. 2015). However, this incidental ammonia cycling would neither yield a net flux of oxidized nitrogen species nor provide electrons for the phototrophic electron transport chain in *Ca.* C. photoalkanotrophicum (oxidation of ammonia by sMMO would be balanced by reduction of hydroxylamine back to ammonia by cytochrome c552 nitrite reductase, resulting only in a net loss of electrons), and so would not therefore be a true case of phototrophic ammonia oxidation.

As with the acquisition of methanotrophy as described here, horizontal gene transfer appears to be a dominant mode of metabolic evolution in the Chloroflexi, particularly involving modular components of high-potential electron transport pathways such as aerobic respiration, denitrification, and phototrophy (Ward et al. 2018a, Crombie and Murrell 2014, Colby et al. 1977, Ward et al. 2018c). Estimates of both relative (via phylogenetic analyses of the antiquity of aerobic respiration in Chloroflexi clades) and absolute (via molecular clock estimates of the radiation of Chloroflexi clades) timing of metabolic diversification in the Chloroflexi suggests that much of this expansion has occurred after the evolution and expansion of aerobic respiration around the Great Oxygenation Event (GOE) ~2.3 billion years ago (Ward et al. 2018a, Fischer et al. 2016). Regardless of the timing of evolutionary innovation in the Chloroflexi, photomethanotrophy as described here must postdate the origin of oxygenic photosynthesis. The initial activation of methane is kinetically inhibited, and is only known to occur without O_2_ in Euryarchaea related to methanogens (Orphan et al. 2001, Knittel and Boetius 2009). All other instances of biological methane oxidation—including photomethanotrophy as described here— rely on O_2_ indicating that this trait most likely postdates the GOE. This includes bacteria which are known to oxidize methane in anaerobic environments via an intra-aerobic pathway driven by production of O_2_ from NO dismutation (Ettwig et al. 2010), a mechanism that requires O_2_-derived substrates and the biochemical capacity for aerobic respiration and aerobic methanotrophy (e.g. Wu et al. 2011). Additionally, the apparent young radiation of phototrophic Chloroflexi (<1 Ga, Shih et al. 2017) suggests that the unique biochemical coupling of methanotrophy and photosynthesis in *Ca.* C. photoalkanotrophicum could not have arisen early in Earth history before the GOE.

**Supplemental Figure 1:** Phylogeny of PufL and PufM protein sequences from a subset of available Chloroflexi genomes, showing the congruence of protein and organismal phylogenies (e.g. Figure 1) in the Chloroflexaceae family, reflecting a history of vertical inheritance of phototrophy genes.

**Supplemental Figure 2:** Phylogeny of A-family heme copper oxidoreductase protein sequences from a subset of available Chloroflexi genomes, showing the congruence of protein and organismal phylogenies (e.g. Figure 1) in the Chloroflexaceae family, reflecting a history of vertical inheritance of respiration genes (though incongruent relationships across the Chloroflexi reflect a history of deeper horizontal gene transfer events).

**Supplemental Figure 3:** Phylogeny of alternative complex III protein sequences from a subset of available Chloroflexi genomes, showing the congruence of protein and organismal phylogenies (e.g. Figure 1) in the Chloroflexaceae family, reflecting a history of vertical inheritance of core electron transport pathway genes (though incongruent relationships across the Chloroflexi reflect a history of deeper horizontal gene transfer events).

**Supplemental Figure 4:** Phylogeny of B-family heme copper oxidoreductase protein sequences from a subset of available Chloroflexi genomes. The sequence from OHK40 is closely related to those of other Chloroflexaceae, but the placement of OHK40 in the phylogeny is incongruent with organismal phylogenies (as part of the *Roseiflexus* clade) no B-family HCO was recovered from the sister taxa *Ca.* Chloroploca asiatica or *Ca.* Virdilinea mediisalina. This may reflect loss in the OHK/Chloroploca/Viridilinea lineage followed by secondary HGT of a B-family HCO by OHK40 from a member of the *Roseiflexus* lineage.

**Supplemental Figure 5:** Phylogeny of cytochrome c552 protein sequences from diverse microbial genomes available on NCBI Genbank and WGS databases, showing that strains closely related to OHK40 lack c552 genes and that the phylogenetic relationships among Chloroflexi c552 proteins are incongruent with organismal relationships, likely reflecting a history of horizontal gene transfer.

**Supplemental Figure 6:** Phylogeny of CO dehydrogenase protein sequences from diverse microbial genomes available on NCBI Genbank and WGS databases, showing that the protein sequence from OHK40 is not closely related to those from other Chloroflexi, and that this gene has likely undergone recent horizontal gene transfer.

**Supplemental Figure 7:** Phylogeny of smmoA beta chain protein sequences from diverse microbial genomes available on NCBI Genbank and WGS databases, showing that the protein sequence from OHK40 is most closely related to distantly related taxa, and that this gene has likely undergone recent horizontal gene transfer. Close relatives to the sequence from OHK40 are similar to those for other smmo subunits, suggesting that the operon underwent HGT intact.

**Supplemental Figure 8:** Phylogeny of smmo regulatory protein B sequences from diverse microbial genomes available on NCBI Genbank and WGS databases, showing that the protein sequence from OHK40 is most closely related to distantly related taxa, and that this gene has likely undergone recent horizontal gene transfer. Close relatives to the sequence from OHK40 are similar to those for other smmo subunits, suggesting that the operon underwent HGT intact.

**Supplemental Figure 9:** Phylogeny of smmo component C protein sequences from diverse microbial genomes available on NCBI Genbank and WGS databases, showing that the protein sequence from OHK40 is most closely related to distantly related taxa, and that this gene has likely undergone recent horizontal gene transfer. Close relatives to the sequence from OHK40 are similar to those for other smmo subunits, suggesting that the operon underwent HGT intact.

**Supplemental Figure 10:** Phylogeny of methyl acetate hydrolase and homologous protein sequences from diverse microbial genomes available on NCBI Genbank and WGS databases, showing the the protein sequence from OHK40 is most closely related to distantly related taxa, and that this gene has likely undergone recent horizontal gene transfer.

**Supplemental Table 1:** Comparison of proteins encoded by *Ca.* Chlorolinea photoalkanotrophicum and *Ca.* Chloroploca asiatica as annotated by RAST.

**Supplemental Table 2:** Comparison of proteins encoded by *Ca.* Chlorolinea photoalkanotrophicum and *Ca.* Viridilinea mediisalina as annotated by RAST.

**Supplemental Table 3:** Proteins encoded by *Ca.* Chlorolinea photoalkanotrophicum but neither of its close relatives *Ca.* Chloroploca asiatica and *Ca.* Viridilinea mediisalina as determined by RAST. Proteins encoded by *Ca.* Chlorolinea photoalkanotrophicum but neither of its close relatives have potentially been recently acquired in this lineage by HGT or by loss in the Chloroploca/Viridilinea lineage. MetaPOAP estimate of False Negative of a single in gene in both relatives is ~0.000025 given their completeness of >99%, providing strong support that these genes are absent from the source genomes. Instances of recent HGT were determined by comparison of topological congruence between protein and organismal phylogenies.

## Notes

#### Summary of Updates

Given uncertainty in the substrate range utilized by this organism, we have revised the manuscript to consider the possibility that this organism oxidizes methane, propane, or other small alkanes.

