## Supplementary figures and images for "Genomic Evidence for Phototrophic Oxidation of Small Alkanes in a Member of the Chloroflexi Phylum"

### Supplemental Figure 1

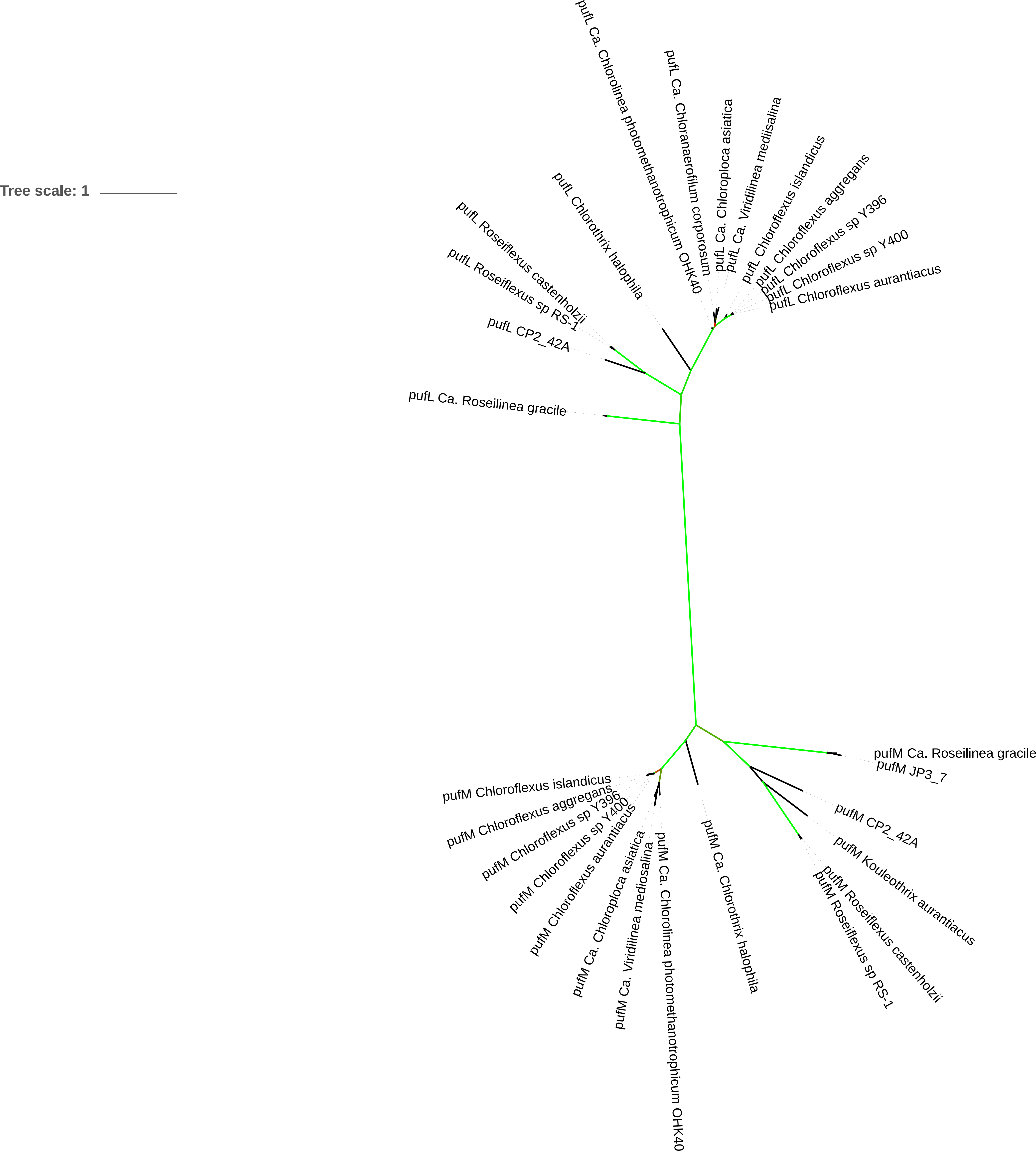

### Supplemental Figure 2

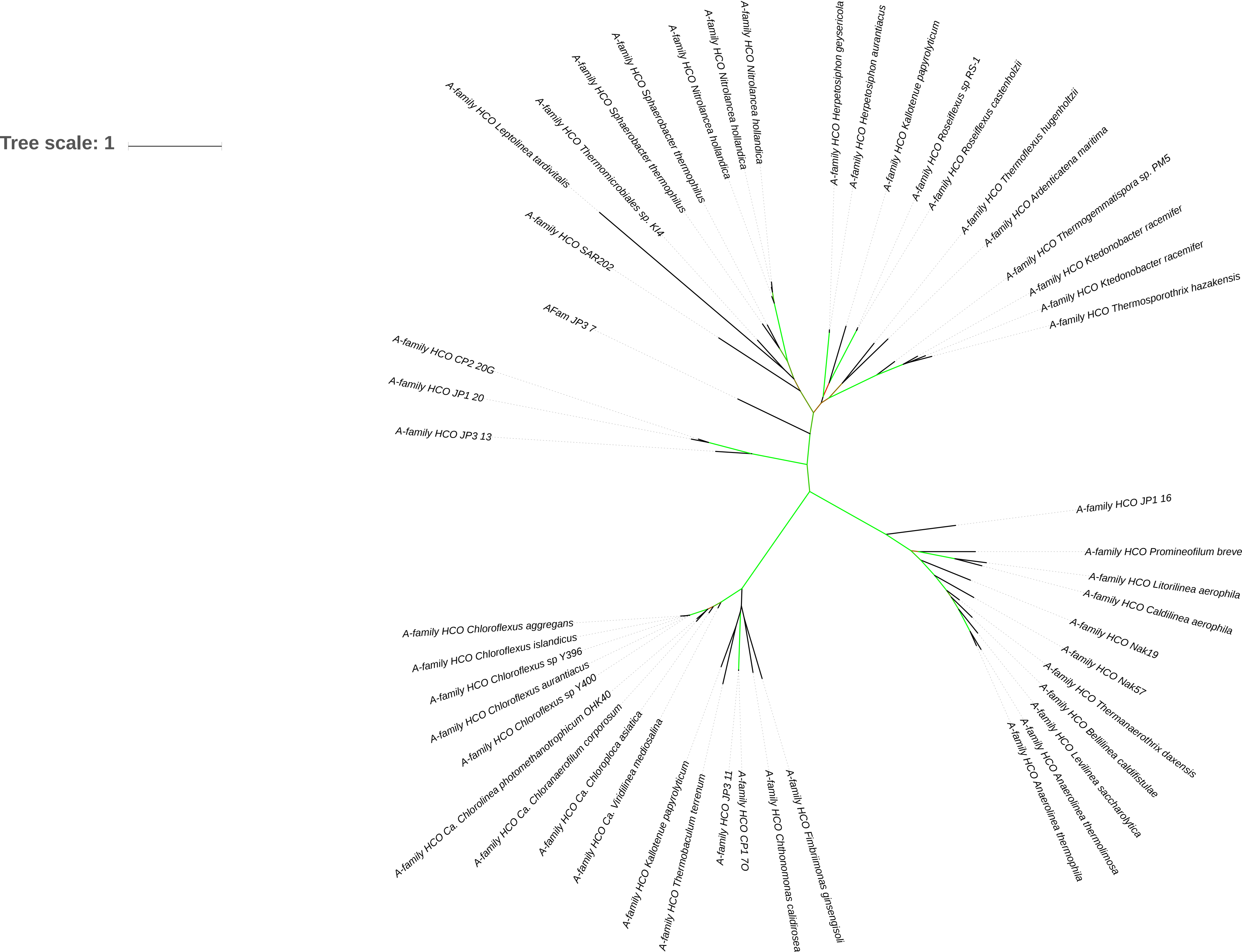

### Supplemental Figure 3

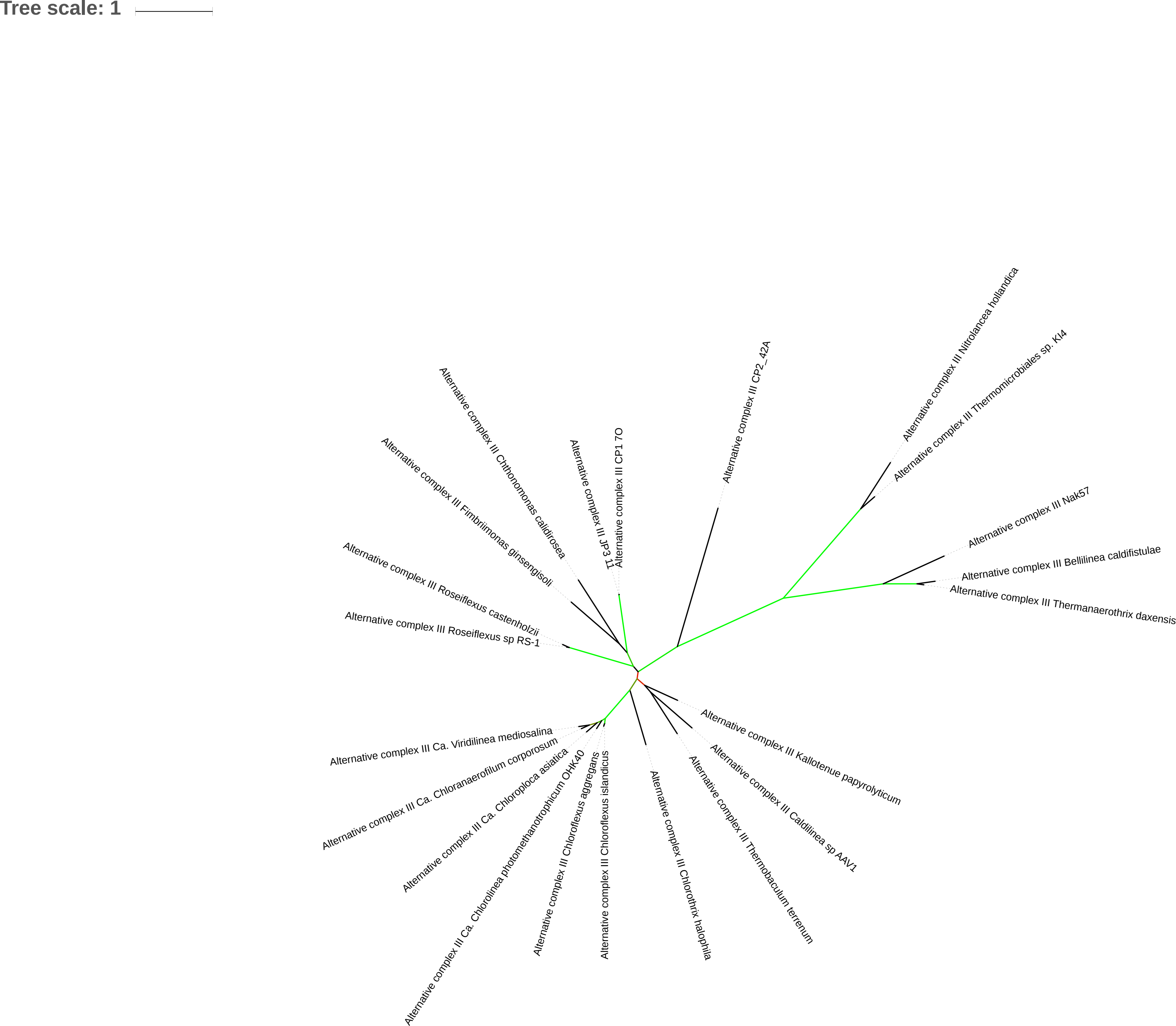

### Supplemental Figure 4

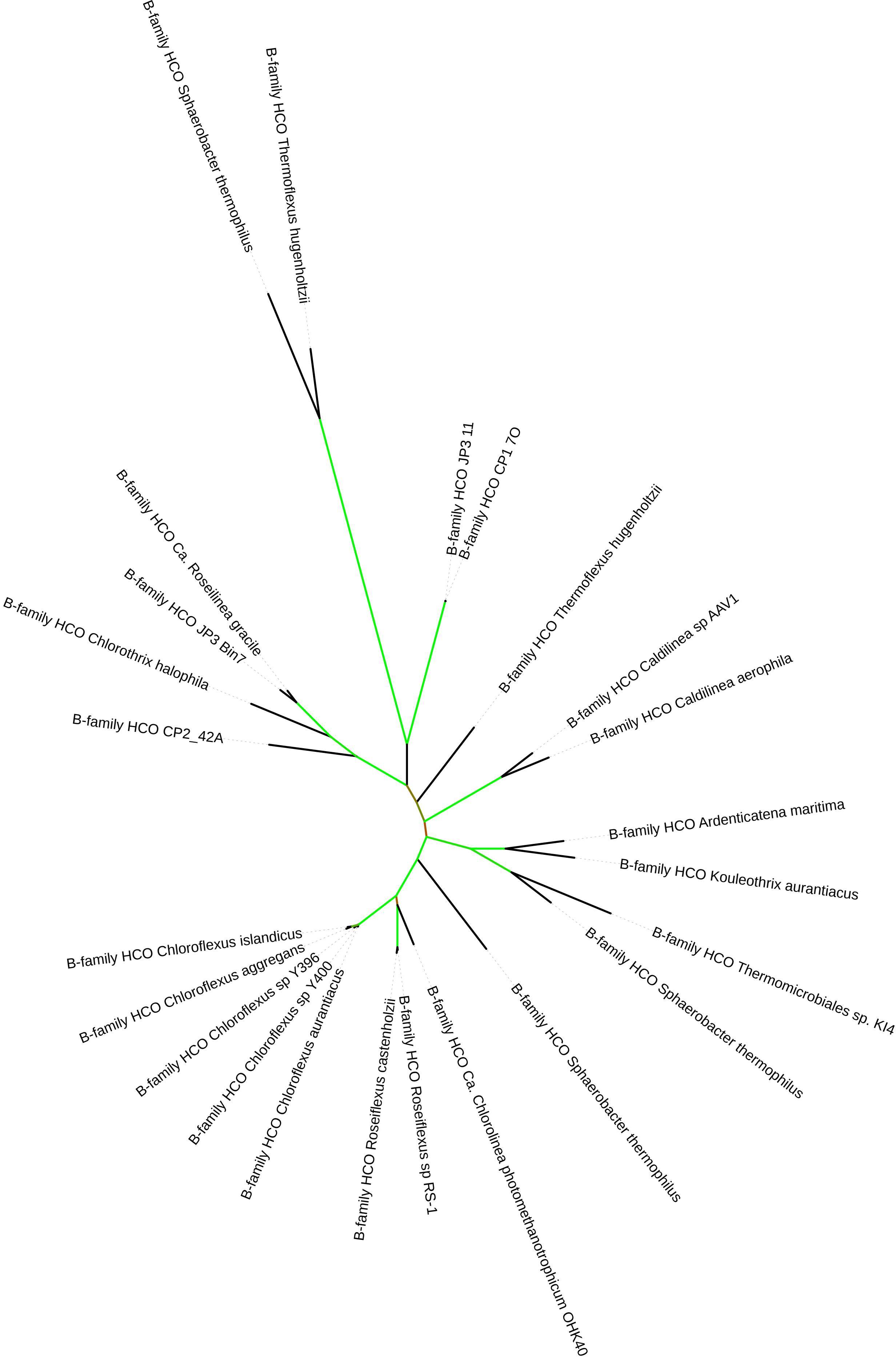

### Supplemental Figure 5

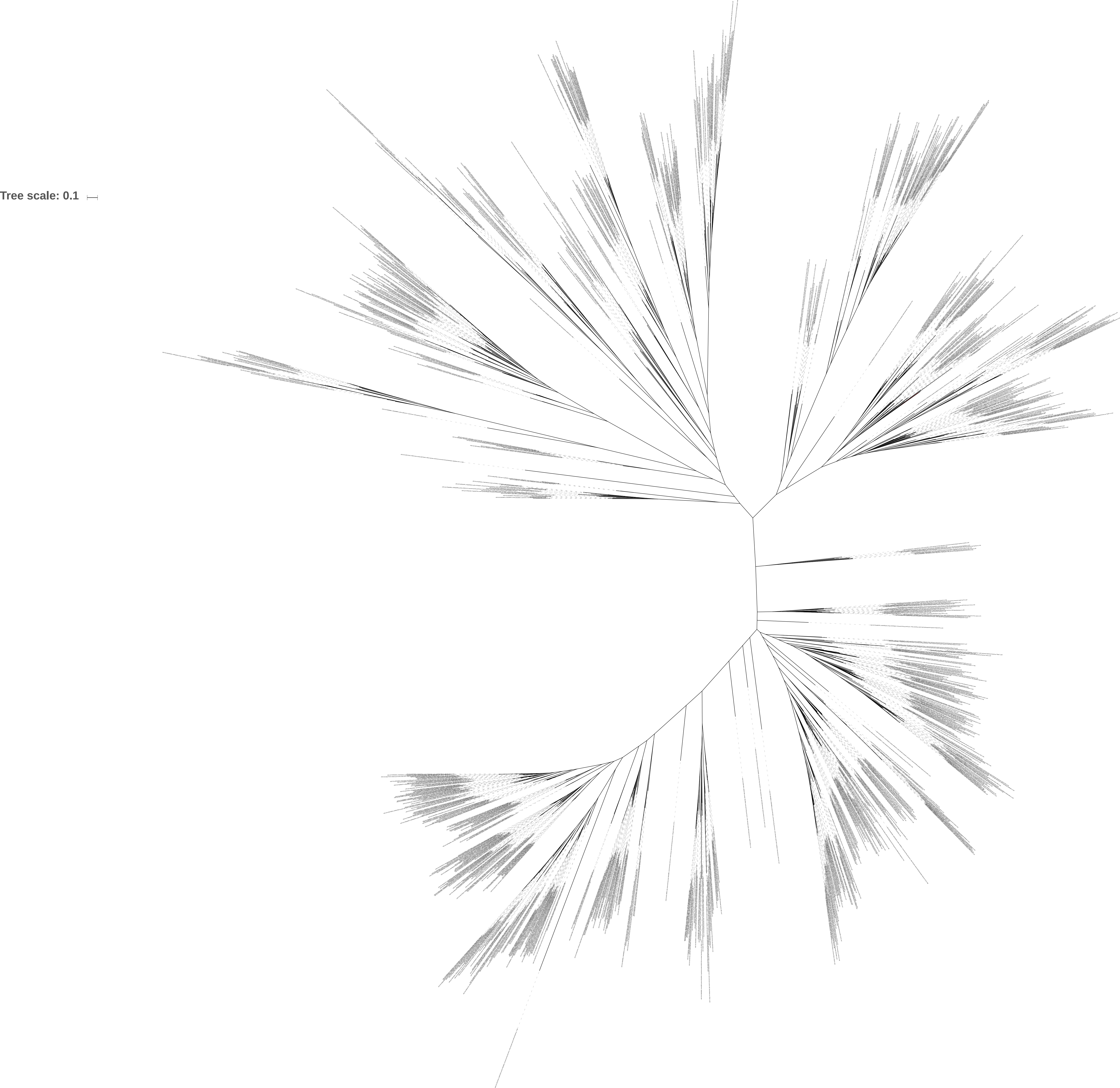

### Supplemental Figure 7

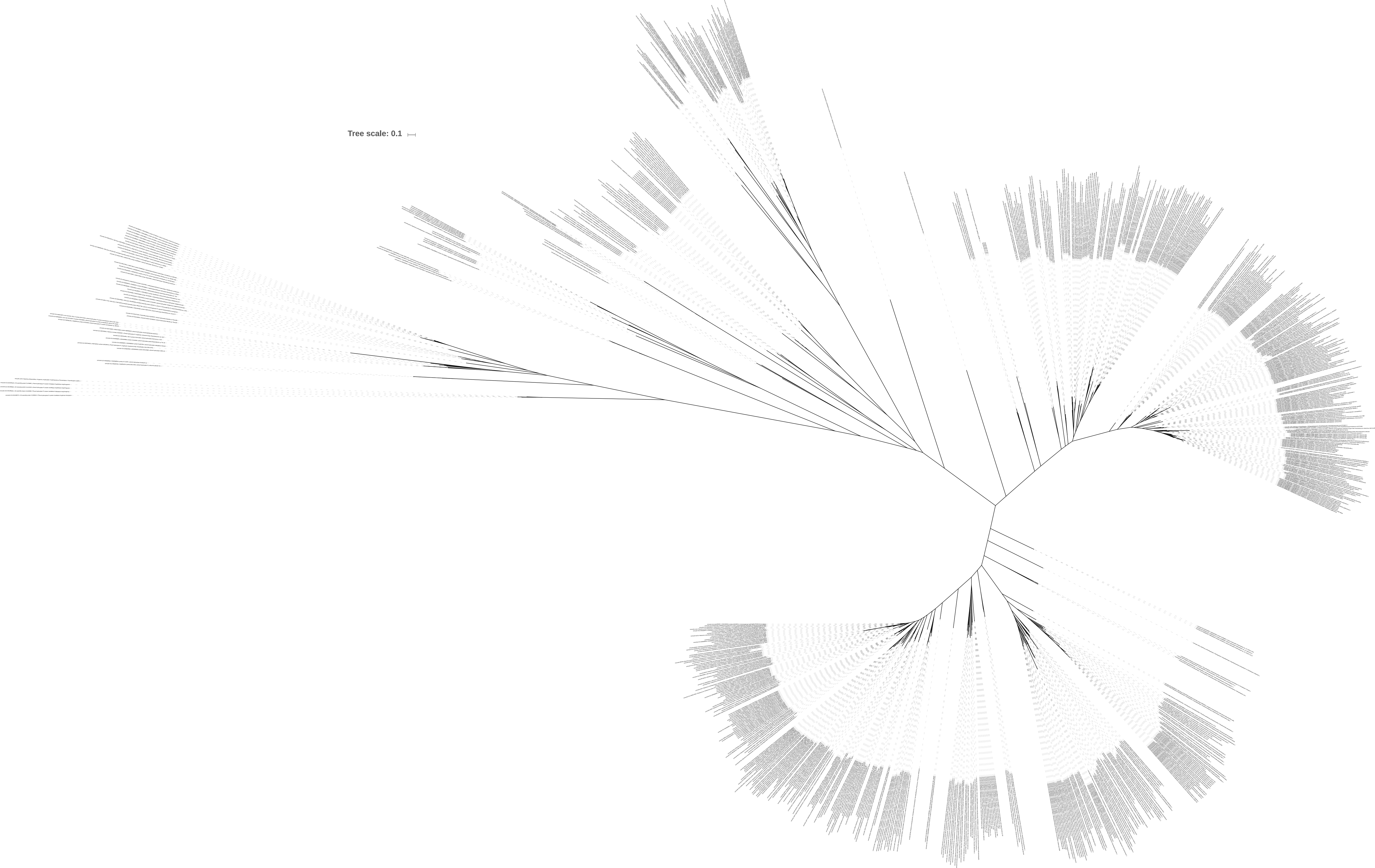

### Supplemental Figure 8

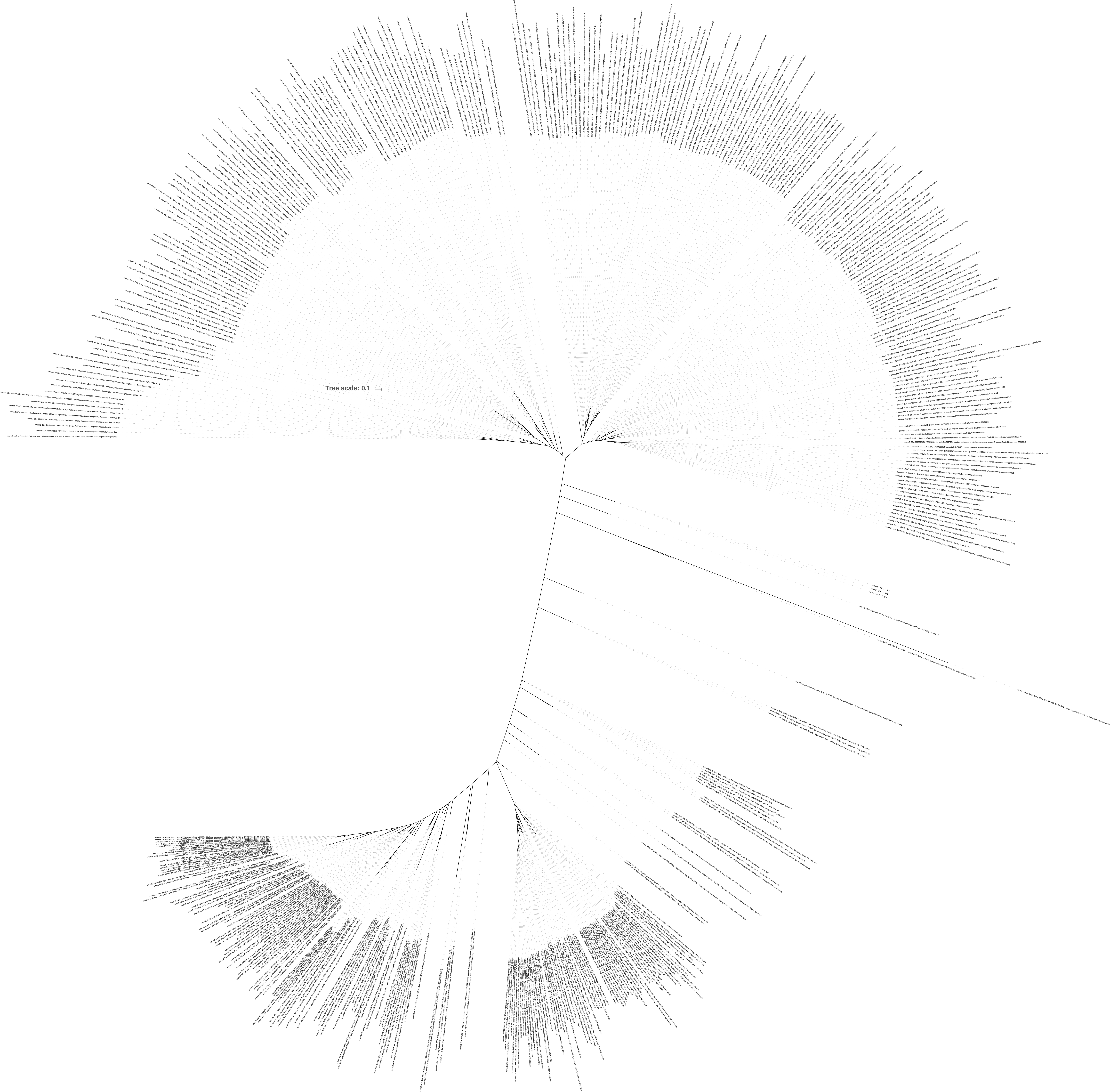

### Supplemental Figure 9

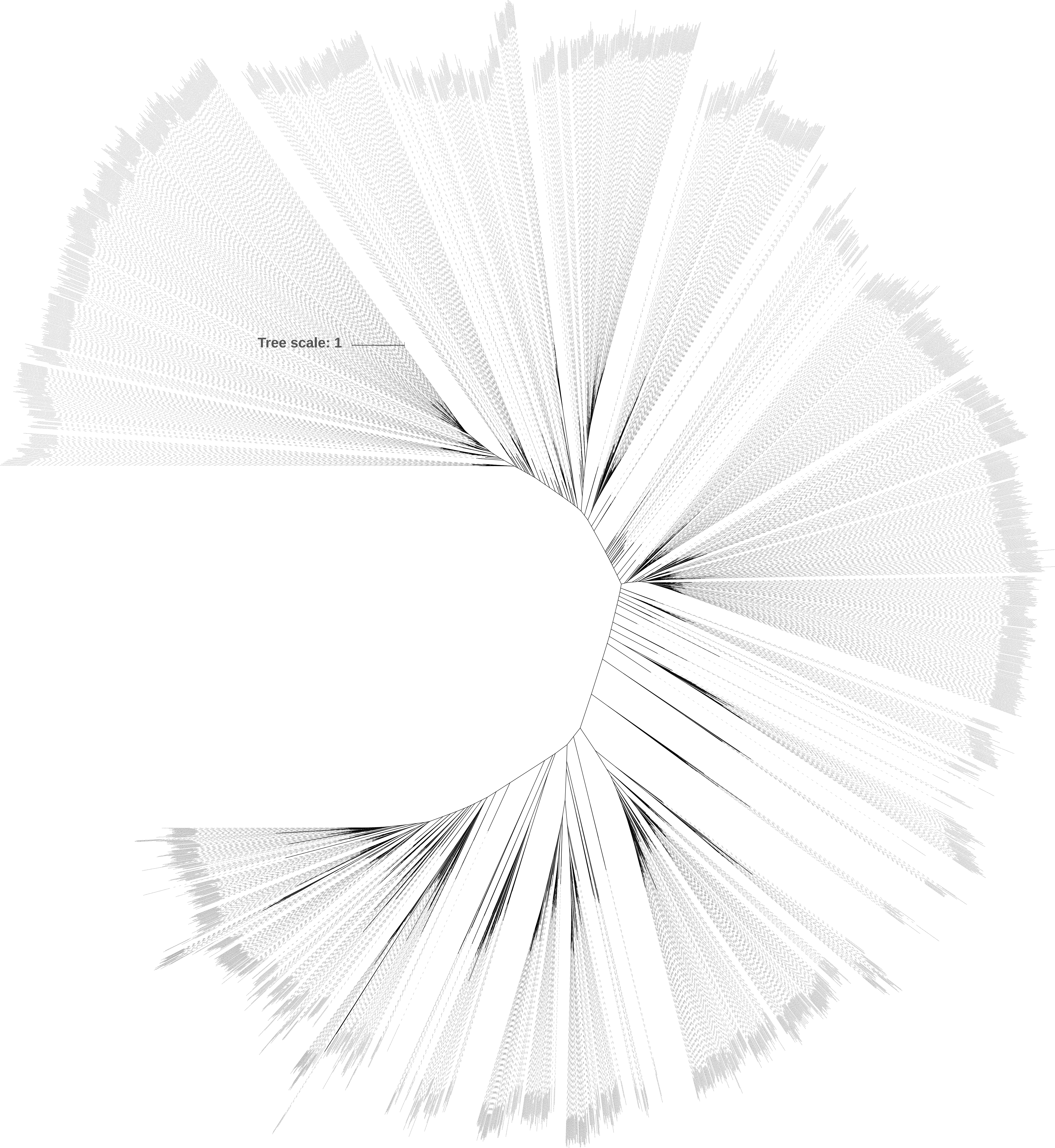

### Supplemental Figure 10

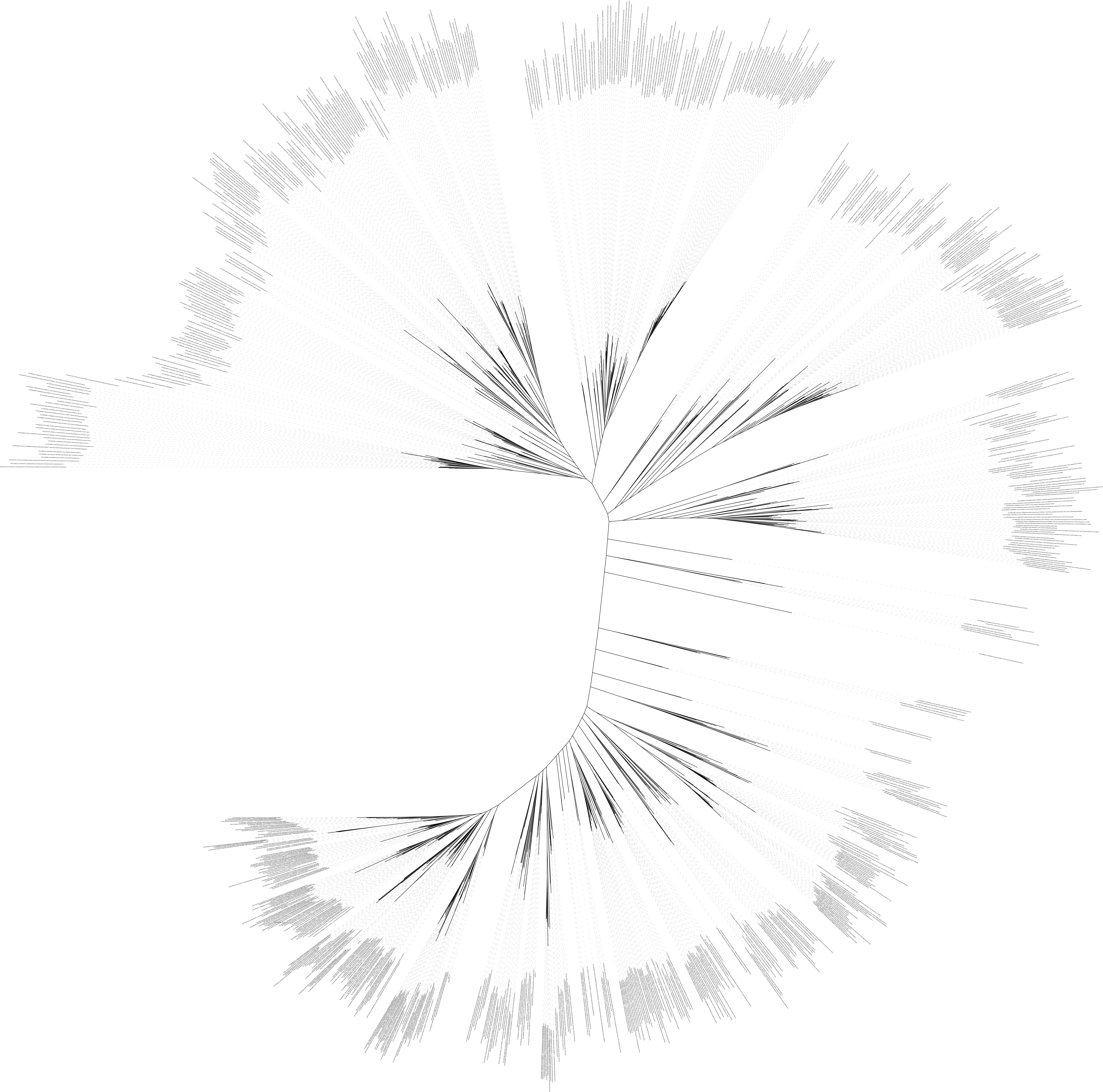
